# Decoding the Dewlap: Multiple signals in females and males of a gliding lizard

**DOI:** 10.1101/2024.08.04.606559

**Authors:** Avantika Deep Sharma, Aravind Sridharan, Kavita Isvaran

## Abstract

Social interactions across taxa are often mediated through multiple signals. Studies examining the maintenance of multiple signals are mostly focused on males and often fail to capture female signalling diversity and strategies. In the recent decade, there has been a surge in the documentation of female signalling, however, our understanding of the functional relevance of multiple signals in females still lags behind. In this study, we examined multiple signals in females of an arboreal gliding lizard, *Draco dussumieri*, and compared them to those in males. We specifically tested the relative role of the backup signal and the multiple receiver hypotheses in the maintenance of multiple signals in both sexes. Female *D.dussumieri* used a variety of signals to socially interact with conspecifics, especially using their dewlap. The signalling repertoire of females was as diverse as that of males, although the relative use of the signals varied. In females, a few signals seem to be maintained by the backup signal hypothesis, with limited support for the multiple receiver hypothesis as well. For males too, both mechanisms appeared to maintain multiple signals. Interestingly, for some signals, the sexes differed in the context in which they used a given signal. Overall, these findings highlight the functional role of multiple signals in females, which can differ from that observed in males. Therefore, traits conventionally considered male-exclusive when also examined in females can provide finer insights into trait function and evolution.

## Introduction

Signalling is a fundamental component of social interaction and mating competition (Rendall et al., 2009; Laidre & Johnstone, 2013). It allows one to assess and convey information to a competitor in a direct contest (Setchell & Jean Wickings, 2005) and to a mate in indirect competition or mate choice (Hill, 2015). Further, in mating contexts, i.e., intra-sexual and inter-sexual interactions, individuals often employ multiple signals (Candolin, 2003). Historically, owing to numerous research biases, studies examining the functional relevance of multiple signals have primarily been carried out in males (T. Clutton-Brock, 2009; Tobias et al., 2012; Driessens et al., 2015). Recent literature suggests female competitive traits and strategies like signalling, ornaments, and aggression are widespread and serve various functions (T. Clutton-Brock, 2007; Stockley & Bro-Jørgensen, 2011; Rosvall, 2011; Rubenstein, 2012; Tobias et al., 2012; T. H. Clutton-Brock & Huchard, 2013; Nolazco et al., 2022). However, studies comparing signal function between the sexes are limited. Uncovering the processes that maintain multiple signals in males and females can be critical to our understanding of different selective pressures that the sexes face and can provide a more comprehensive explanation for signal function and evolution (Driessens et al., 2015; Odom et al., 2016; Riebel et al., 2019; Jones et al., 2021).

Males and females differ in their reproductive roles and biology; hence, their competitive traits and strategies are expected to vary too. The relative energetic investment in reproductive and parental tasks results in differences in the form and intensity of competition and the development of elaborate traits like signals in the sexes (T. Clutton-Brock, 2007, 2009; Stockley & Bro-Jørgensen, 2011). Moreover, in addition to the use of signals in mate competition, a trait they share with males, females’ signalling can extend to competing with individuals of the same sex over other resources such as breeding sites, social rank, and parental care (T. Clutton-Brock, 2009; Rosvall, 2011; Tobias et al., 2012; T. H. Clutton-Brock & Huchard, 2013). Elaborate signalling traits in females, therefore, may serve more diverse functions than those observed in males (Pryke, 2007; Pryke & Griffith, 2007; Odom et al., 2016), potentially affecting signal evolution (Odom et al., 2016; Wilkins et al., 2020; Sierro et al., 2022). Therefore, traits conventionally considered to be male-specific when examined comparatively in both sexes, can provide key insights into trait evolution and sexual dimorphism (Brunton & Li, 2006; Riebel et al., 2019; Wilkins et al., 2020).

A growing number of studies suggest that social interactions are often mediated through multiple signals (Rowe, 1999; Candolin, 2003; Hebets & Papaj, 2005; Bro-Jørgensen, 2010). Based on content, multiple signals could convey either additive information (backup signal hypothesis) (Moller & Pomiankowski, 1993) or distinct messages to multiple receivers (multiple receiver hypothesis) (Marchetti, 1998; Andersson et al., 2002). According to the backup signal hypothesis, multiple signals reinforce the same information, which allows for a more accurate assessment (Moller & Pomiankowski, 1993; Johnstone, 1997a). For example, in the peninsular rock agamas, *Psammophilus dorsalis*, male breeding signals are highly correlated and primarily maintained by female choice (Deodhar & Isvaran, 2018), providing support for the backup signal hypothesis. An alternative explanation proposes that the multiple signals are maintained as they are directed towards different receivers (the multiple receiver hypothesis). Different receivers might pay attention to different kinds of information or different aspects of the signalling individual (Marchetti, 1998; Andersson et al., 2002). For example, in several species, males may display using signals with multiple components, in which one component conveys mating ability to females while another component signals competitive ability to other males (Andersson et al., 2002; Loyau et al., 2005; Driessens et al., 2014).

Further, depending on the sexual identity of the signalling individual, the same signal could play a different role. In multiple bird species, the same traits have been documented to indicate a different quality in the sexes. For instance, in common yellowthroats, *Geothlypis trichas*, the brightness of the bib signals fecundity in males but indicates poor health in females (Dunn et al., 2010; Freeman-Gallant et al., 2014). Similarly, in northern cardinals, *Cardinalis cardinalis*, the facial pattern correlates with high reproductive success in males (Jawor & Breitwisch, 2004), while in females, the same facial marks predict intra-sexual aggression (Jawor et al., 2004). These differences could arise because of the different selective pressures the signal might be under in each sex (Harrison & Poe, 2012; Odom et al., 2016; Webb et al., 2016). This difference might also extend to a differential mechanism that maintains multiple signals in each sex. For instance, due to the higher cost of signalling traits and to ensure efficient transmission of information, female signals might be more correlated (reinforcing) than male signals (Candolin, 2003; T. Clutton-Brock, 2007; Webb et al., 2016). Conversely, with the diverse functions female signals perform in both breeding and non-breeding contexts, multiple signals might be maintained to convey different information to different receivers (Tobias et al., 2012; Odom et al., 2016). As a result, the function of multiple signals derived from male-focused studies cannot be extrapolated to female signalling traits.

In this work, we tested the relative importance of the multiple receivers hypothesis (Andersson et al., 2002) and backup signals hypothesis (Moller & Pomiankowski, 1993) in maintaining multiple signals in both sexes in a gliding lizard, *Draco dussumieri*. It is a medium-sized arboreal agamid endemic to India. Male *D.dussumieri* have been documented to use multiple visual signals for intra-specific communication; however, similar information for females is lacking (John, 1962, 1966). There has been no record of olfactory or acoustic communication, thus, allowing the examination of the complete signalling repertoire of the species. We assessed what maintains multiple signals in both sexes by examining the associations of the signals with two social contexts, namely intra-sexual and inter-sexual interactions. Strongly correlated signals associated with the same social context would indicate support for the back-up signal hypothesis (Bro-Jørgensen & Dabelsteen, 2008). Conversely, if the signals are uncorrelated, with some signals performed in intra-sexual interactions while others in inter-sexual interactions, it would imply that multiple signals are intended for multiple receivers (Andersson et al., 2002; Candolin, 2003).

## Methods

### Study System

This study was conducted from January 2022 to April 2022 in three different privately owned areca nut plantations in the Agumbe landscape (13°50′ N, 75°09′ E; 560 m above sea level) in Karnataka, India.

*Draco dussumieri* (Indian gliding lizard) is a medium-sized gliding agamid endemic to India. It is a strictly arboreal lizard, with only the females coming down the tree to lay eggs on the ground (John, 1962, 1966). The lizard has a lateral wing membrane on each side of its body called a patagium, which is formed by the skin supported by ribs. It enables the lizard to glide among the trees, a unique feature of this genus. Additionally, the species also has a gular appendage, or dewlap, which is extendible. It acts as an organ for communication and social display and is larger in males than females (John, 1962, 1966; Mori & Hikida, 1993, 1994; Ferguson, 1977; Hairston, 1957; Harrison & Poe, 2012; Klomp et al., 2016, 2017). The species is reported to breed from February to April (John, 1966); however, subsequent information about its life history and development is deficient.

### Data Collection

For individual identification, we caught lizards using a long (30 ft) telescopic pole. They were induced to glide down to the ground by nudging with the end of the pole (Mori & Hikida, 1993; Sreekar et al., 2013). Using a white, non-toxic permanent marker, we gave each individual a unique number (Sreekar et al., 2013). Morphometric data, which included weight, snout-vent length, head width, dewlap length, and tail length, was also collected for all the caught individuals using an electronic weighing machine and vernier caliper. Handling time lasted a maximum of 10 minutes per individual. Lizards were released back on the same tree that they were captured from. All the animal handling and behavioural sampling complied with the guidelines of the Institutional Animal Ethics Committee constituted by the National Centre for Biological Sciences [NCBS-IAE-2021/ 06(N)].

Behavioural data was collected by carrying out focal animal sampling (Altman, 1974). We recorded behaviour of both marked and unmarked individuals selected opportunistically. However, effort was made to collect behavioural data from each marked individual at least once over the study period. During the sampling, an individual’s behaviour was recorded continuously for 20 minutes using a Panasonic camcorder. If the individual disappeared within five minutes of observation and did not return, the observation was discarded. We also recorded the duration of time male and female conspecifics were present within 5 metres of the focal individual during a focal animal sampling session. The distance of 5m was chosen based on pilot observations, as most dyadic interactions were seen within this distance. Additionally, information related to the weather was also noted. Based on the visual estimate, if the cloud cover was more than 80%, the weather was considered cloudy; otherwise, it was classified as sunny.

Video recordings were transcribed using the software BORIS (Behavioural Observation Research Interactive Software) version 7.12.2 (Friard & Gamba, 2016). While transcribing, short-duration behavioural events and long-duration behavioural states were identified and uniquely coded. Based on previous literature (John, 1962, 1966; Hairston, 1957; Mori & Hikida, 1993, 1994) and personal observations we identified key signalling behaviours like dewlap extension, tail raise, patagium extension etc. Other behaviours not involved in display like locomotion, resting, foraging etc. were also identified. For a detailed list of behaviours and their definitions, see Supplementary Table 1. The time for which the focal individual moved out of the frame was removed from the observation period. The duration for which a conspecific male or female was within 5m of the focal individual was also coded and logged.

### Statistical analysis

All the analyses were carried out in R version 4.13 (R Core Team, 2022). The behaviours were quantified by calculating the frequency (reported as count per 300 seconds) for events and the proportion (time spent in a specific behaviour divided by total time sampled) of time spent for states. The duration for which a male or female conspecific was present within 5 m was also quantified for each focal observation as proportion (time for which conspecific was present divided by total time).

To check for correlation in signals, we performed a principal components analysis using the R package FactoMineR (Jose & Husson, 2008) separately for each sex. If the signals load strongly on the same axis, they might be covarying or backup signals (Bro-Jørgensen & Dabelsteen, 2008).

To check for the intended receiver of the signals, we analysed the relationship between the signals and social context. Each male and female signal was modelled as a function of the proportion of time a male and female conspecific was present within a 5m radius. For event behaviours (represented as counts), GLMs were run using the package pscl (Jackman, 2020). To correct for zero inflation and overdispersion, a two-step hurdle model was used. The time for which the individual was visible was included as an offset term to account for variability in the sampling effort. Beta regression was used for analysing state behaviours (proportion of time), and models were fit using the R package betareg (Cribari-Neto & Zeileis, 2010). Since the response variable had some zeros, to avoid violating the assumptions of beta regression, we added an offset value of 0.01.

For all the models permutation tests were run and Type S error (Gelman & Weakliem, 2009) and 95% bootstrapped confidence interval on the model estimate were also calculated. This was done to increase our confidence in the models and to make inferences based on a composite of results rather than just based on p-values.

## Results

We caught and uniquely identified forty-two *D.dussumieri* individuals; of these, 23 were males and 18 were females. Hundred and sixty-nine focal observations were carried out with repeated measures on 32 (12 females, 20 males) of the marked and 35 (10 females, 25 males) unmarked individuals over four months. Out of the hundred and sixty-nine observations, eighty-nine had a conspecific presence within a 5m radius of the focal individual. Focal observations revealed that females engage in multiple signalling behaviours. We observed 10 distinct signals, including dewlap extension, dewlap display, patagium extension, arched back, tail flick, etc. (Supplementary Table 1). Several of these behaviours have been reported in other lizard groups as well (Radder et al., 2006; Patankar et al., 2013; Driessens et al., 2014; Ramos & Peters, 2016; Reedy et al., 2017; Ranade, 2022). The signalling repertoire of females was similar to that of males, except for a few sex-specific signals (Supplementary Table 2). However, the sexes differed in the relative use of the signals (Supplementary Table 2). For instance, males but not females commonly displayed dewlap flicker (occurring in 56% and 5% of male and female focal observations, respectively). Arching of the back was observed four times more frequently in females (29%) than in males (7%) (Supplementary Table 2). These multiple behaviours were observed during specific contexts thereby prompting a need to understand the correlation of signals. Rare behaviours (observed in fewer than 10% of focal observations) were not included in subsequent analyses.

**Figure 1:**
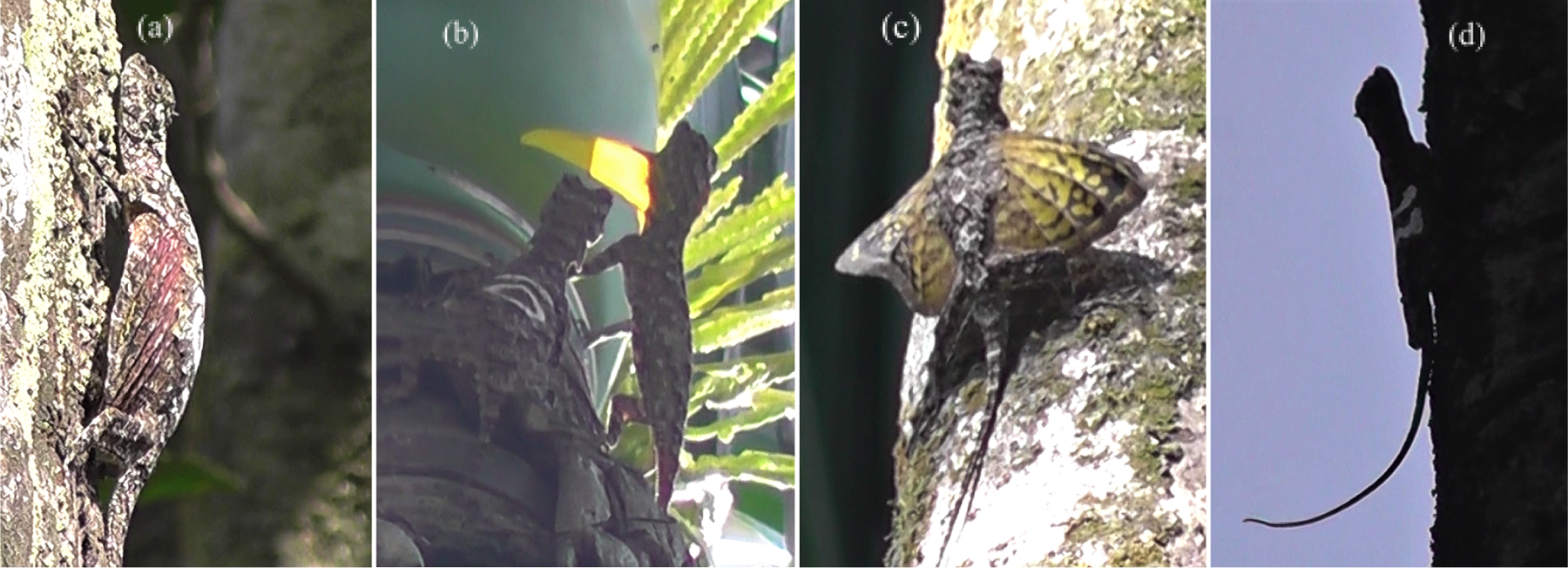
Photographic representation of some signals (a) A female arching its back, (b) A male performing dewlap display while circling around a female, (c) A male extending its patagium, and (d) Tail raise

### Correlation in signals

A PCA was performed separately for each sex to check for correlation in signals. For females, the PC1 axis explained 44.50 % of the variation and PC2 20.76% of the total variation (Figure 2). Signals showed some covariation with relatively high loadings of patagium extension and arched back on PC1.

**Figure 2:**
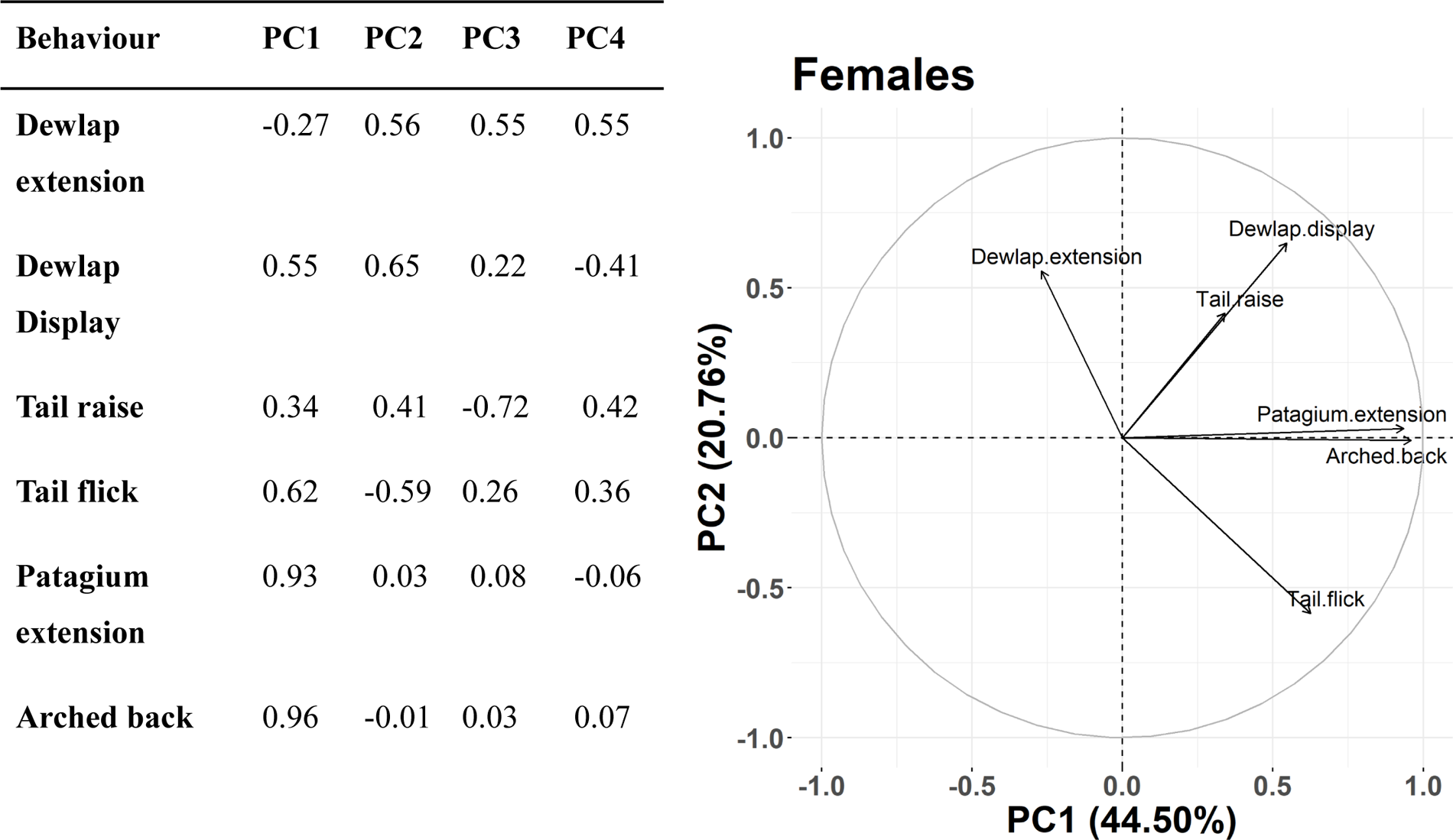
Principal Component Analysis (PCA) on the signals of females. The table (left) shows the loading of signals on each axis. The graph (right) shows covarying behaviours. Each line represents one signal.

For males, the PC1 explained 32.98% of the variation and the PC2 explained 21.16% of the total variation. Dewlap extension and slow dewlap extension had relatively high loading on PC1, while head bob and dewlap display loaded strongly on PC2. The other signals did not load strongly on any particular axis. (Figure 3). Further, for both females and males, since signals were not strongly correlated, a composite variable could not be used in the analysis of social context, and each signal was analysed separately.

**Figure 3:**
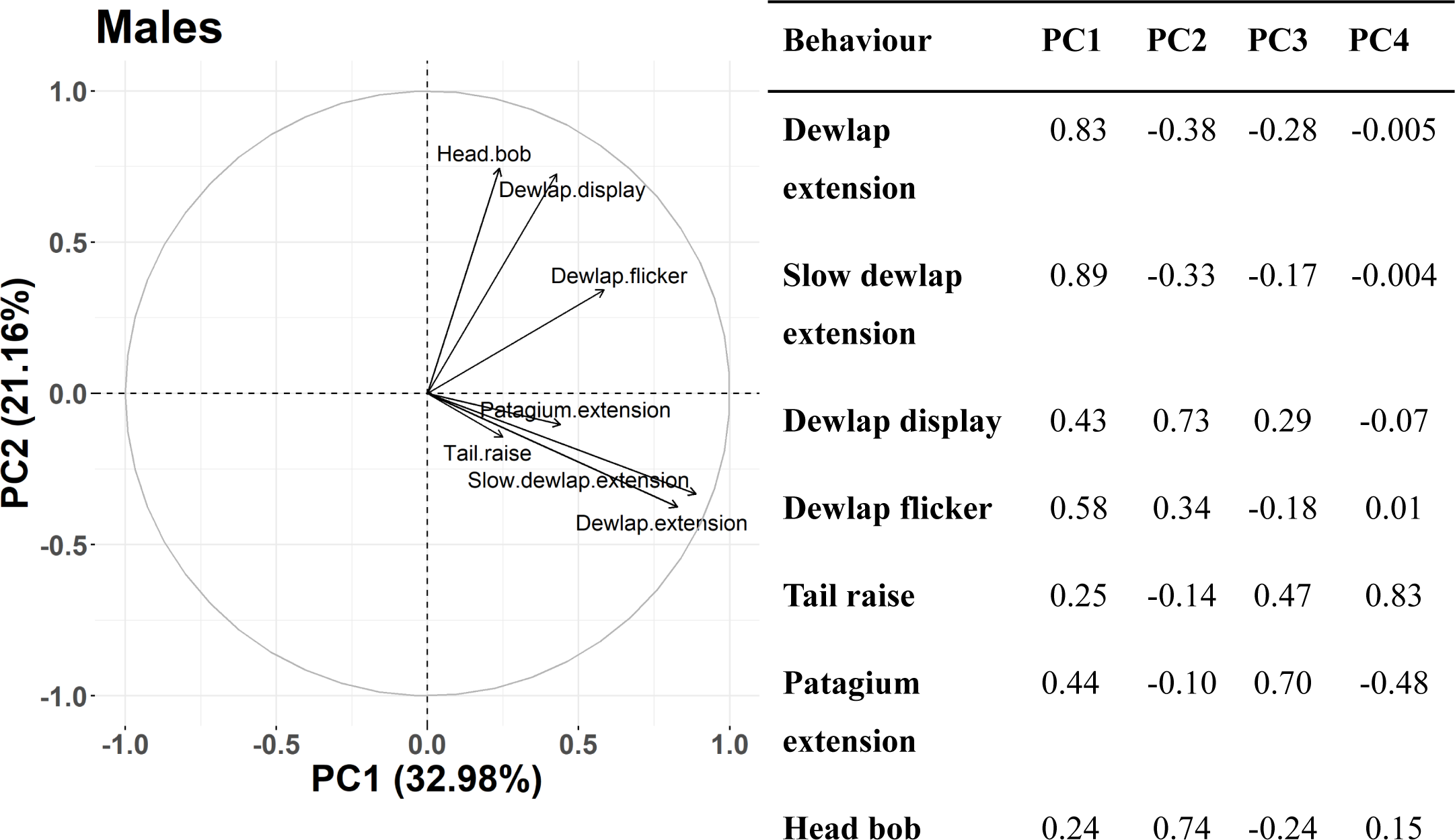
Principal Component Analysis (PCA) on signals of males. The table (right) shows the loading of signals on each axis. The graph (left) shows covarying behaviours. Each line represents one signal.

### Association of signals with social contexts

To check for the association of signals with different social contexts (intra-sexual and inter-sexual), signals (counts for events and proportions for states) were modelled as a function of the proportion of time a male or female conspecific was present within a 5m radius of the focal individual.

In female lizards, some signals were found to be positively associated with the proportion of time a female conspecific was in the vicinity [e.g. dewlap display (χ 2= 21.98, df=1, p<0.01), patagium extension (χ 2= 2.79, df=1, p=0.06) and dewlap extension (χ 2= 1.89, df=1, p=0.09)] while some signals [e.g. patagium extension (χ 2= 2.68, df=1, p=0.10), arched back (χ 2= 2.66, df=1, p=0.10) ] were weakly correlated with the proportion of time a male conspecific was in the vicinity (Table 1, Figure 4). These associations had substantially positive but variable effect sizes and therefore required validation. Based on a very low Type S error value, (error of concluding that the estimate is positive when the true value is negative) and 95% bootstrapped confidence interval of the model estimate we infer that in case of dewlap extension, arched back and patagium extension there is sufficient evidence that the signals are positively associated with male and female conspecifics, but the wide confidence intervals indicate that they require further exploration (Table 1). For instance, the increase in the use of arched back in the presence of conspecific females has a substantially positive 95% bootstrapped confidence interval (-0.02, 1.45) and the probability of erroneously concluding that this estimate is positive when it is actually negative is extremely low (Type S error = 0.0004), suggesting a positive association. Dewlap flicker had very few observations (less than 10% of focal observations) and, therefore, was not included in the analysis.

The results for male signals suggest that some signals [e.g. dewlap display (χ 2= 15.22, df=1, p<0.01), p<0.01), dewlap flicker (χ 2= 5.65, df=1, p<0.01)] were positively associated with the presence of female conspecific, while some signals [e.g. patagium extension (χ 2= 8.35, df=1, p<0.01)] were positively associated with the presence of male conspecific (Table 2, Figure 5). The behaviour turn and circle seemed to be a multi-component signal with dewlap display and tail raise being performed together, and therefore, was excluded from the analysis. Weather did not seem to affect any behaviour in a statistically detectable manner (Tables 1 and 2).

**Table 1:** Association of female signals with the proportion of time male and female conspecifics were in the vicinity. Weather which could affect signalling behaviour, was also included in the model. Results from GLM and betaregression include estimate, standard error, and likelihood ratio tests. Type-S error, permutation test p-value and bootstrapped 95% CI are also reported. Statistically significant associations have been represented in bold. (N=55, df=1)

|  | Dewlap extension |  |  |  |  |  |  |
| --- | --- | --- | --- | --- | --- | --- | --- |
| | Estimate | SE | $\chi^2$ | p | Type-S error | p from permutation test | Bootstrapped 95% CI |
| Intercept | -2.87 | 0.43 |  |  |  |  |  |
| Male conspecific | -0.05 | 0.41 | 0.07 | 0.90 | 0.36 | 0.85 | -0.93, 0.59 |
| **Female conspecific** | **0.72** | **0.43** | **1.89** | **0.09** | **0.0004** | **0.13** | **-0.07, 1.91** |
| Weather:Sunny | -0.17 | 0.37 | 0.02 | 0.63 |  |  |  |

| Tail raise |  |  |  |  |  |  |  |
| --- | --- | --- | --- | --- | --- | --- | --- |
| | Estimate | SE | $\chi^2$ | p | Type-S error | p from permutation test | Bootstrapped 95% CI |
| Intercept | -3.06 | 0.32 |  |  |  |  |  |
| Male conspecific | -0.01 | 0.31 | 0.001 | 0.97 | 0.46 | 0.94 | -0.43, 0.40 |
| Female conspecific | 0.27 | 0.36 | 0.55 | 0.46 | 0.03 | 0.26 | -0.24, 1.22 |
| Weather:Sunny | -0.04 | 0.28 | 0.02 | 0.88 |  |  |  |
| Dewlap Display |  |  |  |  |  |  |  |
| | Estimate | SE | $\chi^2$ | p | Type-S error | p from permutation test | Bootstrapped 95% CI |
| Intercept | -4.60 | 0.11 |  |  |  |  |  |
| Male conspecific | 0.15 | 0.11 | 1.79 | 0.18 | 0.002 | 0.18 | -0.06, 0.38 |
| **Female conspecific** | **0.67** | **0.11** | **21.98** | **<0.01 \*** | **~0** | **0** | **0.16, 1.21** |
| Weather:Sunny | -0.02 | 0.09 | 0.04 | 0.85 |  |  |  |
| Patagium extension |  |  |  |  |  |  |  |
| | Estimate | SE | $\chi^2$ | p | Type-S error | p from permutation test | Bootstrapped 95% CI |
| Intercept | -3.41 | 0.32 |  |  |  |  |  |
| **Male conspecific** | **0.48** | **0.30** | **2.68** | **0.10** | **0.0005** | **0.04** | **0.11, 0.95** |
| **Female conspecific** | **0.61** | **0.33** | **2.79** | **0.06** | **0.0001** | **0.01** | **0.01, 1.64** |
| Weather:Sunny | 0.07 | 0.27 | 0.07 | 0.79 |  |  |  |

| Arched back |  |  |  |  |  |  |  |
| --- | --- | --- | --- | --- | --- | --- | --- |
| | Estimate | SE | $\chi^2$ | p | Type-S<br>error | p from<br>permutation<br>test | Bootstrapped<br>95% CI |
| Intercept | -3.54 | 0.31 |  |  |  |  |  |
| Male conspecific | **0.48** | **0.29** | **2.66** | **0.10** | **0.0004** | **0.03** | **0.08, 0.92** |
| Female conspecific | **0.53** | **0.32** | **2.24** | **0.10** | **0.0004** | **0.04** | **-0.02, 1.45** |
| Weather:Sunny | -0.06 | 0.26 | 0.06 | 0.81 |  |  |  |
| Tail flick |  |  |  |  |  |  |  |
| | Estimate | SE | $\chi^2$ | p | Type-S<br>error | p from<br>permutation<br>test | Bootstrapped<br>95% CI |
| Intercept | -4.47 | 0.12 |  |  |  |  |  |
| Male conspecific | 0.15 | 0.12 | 1.47 | 0.23 | 0.002 | 0.17 | 0.01, 0.03 |
| Female conspecific | -0.07 | 0.15 | 0.24 | 0.63 | 0.10 | 0.61 | -0.19, 0.02 |
| Weather:Sunny | -0.04 | 0.11 | 0.14 | 0.71 |  |  |  |

**Figure 4:**
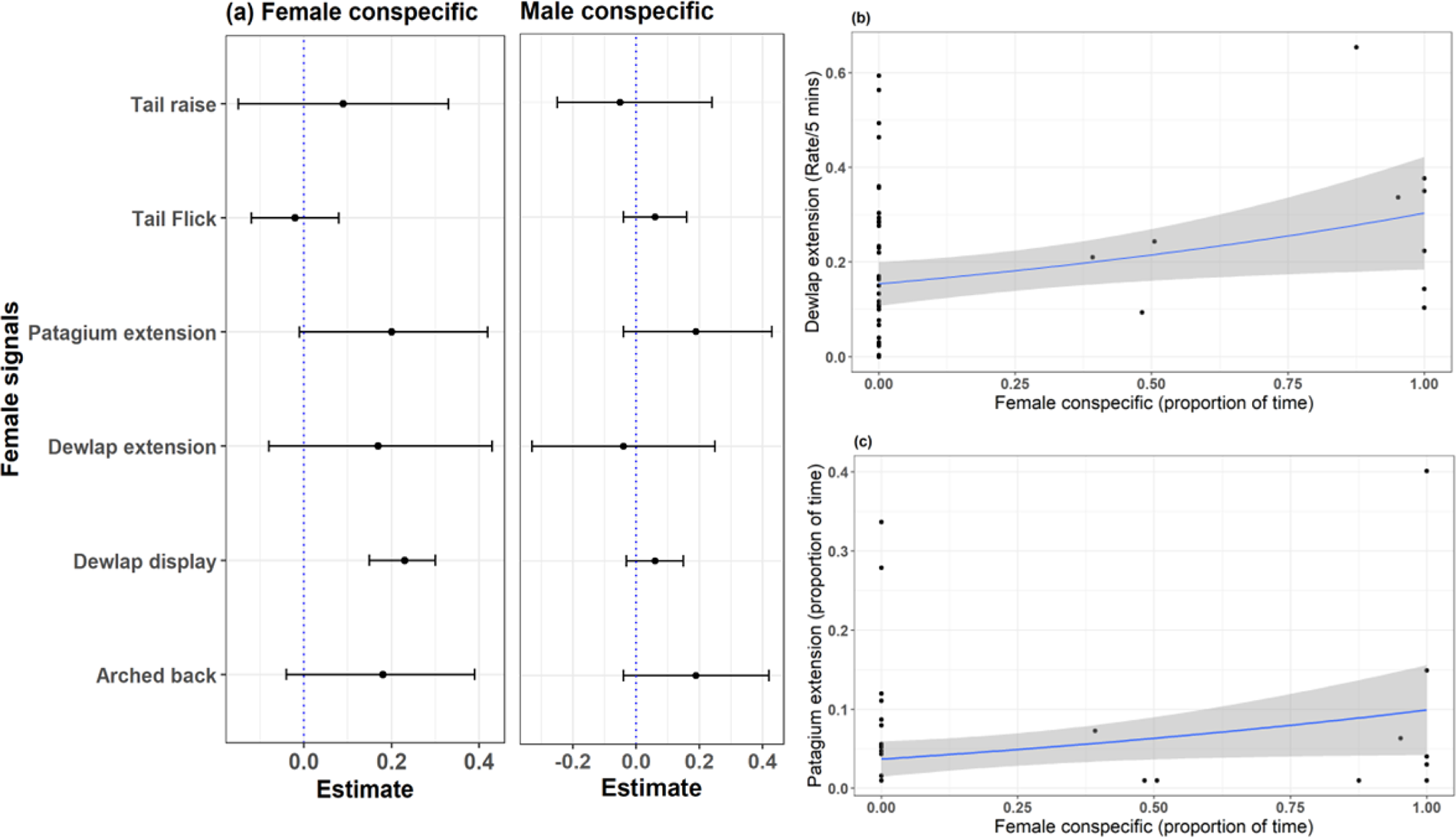
Association of female signals with the presence of male and female conspecifics. (a) Summary of standardised model predictions as represented by effect size and 95 percent confidence interval. (b) and (c) The model prediction represented by the regression line shows the relationship of female signals with female audiences. Dewlap extension (b) and patagium extension (c) were positively related to the proportion of time a female conspecific was in the vicinity. Black dots represent actual data points.

**Table 2:** Association of male signals with the presence of male and female conspecifics. Weather which could affect the signalling behaviour, was also included in the model. Results from GLM and betaregression include estimate, standard error, and likelihood ratio tests. Type-S error, permutation test p-value and bootstrapped 95% CI are also reported. Statistically significant associations have been represented in bold. (N=114, df=1)

| **Dewlap extension** |  |  |  |  |  |  |  |
| --- | --- | --- | --- | --- | --- | --- | --- |
| | Estimate | SE | $\chi^2$ | p | Type-S error | p from permutation test | Bootstrapped 95% CI |
| Intercept | -2.90 | 0.16 |  |  |  |  |  |
| Male conspecific | 0.17 | 0.22 | 0.37 | 0.44 | 0.03 | 0.54 | -0.42, 0.69 |
| Female conspecific | 0.20 | 0.23 | 1.20 | 0.38 | 0.02 | 0.45 | -0.33, 0.88 |
| Weather: Sunny | -0.04 | 0.16 | 0.06 | 0.76 |  |  |  |
| **Slow dewlap extension** |  |  |  |  |  |  |  |
| | Estimate | SE | $\chi^2$ | p | Type-S error | p from permutation test | Bootstrapped 95% CI |
| Intercept | -4.02 | 0.16 |  |  |  |  |  |
| **Male conspecific** | **0.45** | **0.21** | **4.0** | **0.03** | **0.00003** | **0.13** | **-0.12, 1.01** |
| Female conspecific | 0.33 | 0.22 | 2.54 | 0.12 | 0.0008 | 0.23 | -0.16, 0.93 |
| Weather:Sunny | -0.14 | 0.15 | 0.66 | 0.38 |  |  |  |
| **Dewlap flicker** |  |  |  |  |  |  |  |
| | Estimate | SE | $\chi^2$ | p | Type-S error | p from permutation test | Bootstrapped 95% CI |
| Intercept | -6.20 | 0.83 |  |  |  |  |  |
| Male conspecific | 0.62 | 0.64 | 0.68 | 0.33 | 0.01 | 0.48 | -0.89, 1.87 |
| **Female conspecific** | **1.55** | **0.62** | **5.65** | **0.01\*** | **~0** | **0.09** | **0.07, 2.99** |
| Weather:Sunny | -0.82 | 0.47 | 2.83 | 0.08 |  |  |  |

| Head bob |  |  |  |  |  |  |  |
| --- | --- | --- | --- | --- | --- | --- | --- |
| | Estimate | SE | $\chi^2$ | p | Type-S error | p from permutation test | Bootstrapped 95% CI |
| Intercept | -5.27 | 0.96 |  |  |  |  |  |
| Female conspecific | 0.39 | 1.11 | 0.12 | 0.72 | 0.16 | 0.73 | -3.77, 2.88 |
| Weather:Sunny | -0.25 | 0.63 | 0.16 | 0.69 |  |  |  |

| Tail raise |  |  |  |  |  |  |  |
| --- | --- | --- | --- | --- | --- | --- | --- |
| | Estimate | SE | $\chi^2$ | p | Type-S error | p from permutation test | Bootstrapped 95% CI |
| Intercept | -3.58 | 0.20 |  |  |  |  |  |
| Male conspecific | 0.28 | 0.22 | 1.51 | 0.21 | 0.002 | 0.05 | -0.02, 0.64 |
| Female conspecific | 0.09 | 0.25 | 0.14 | 0.71 | 0.16 | 0.57 | -0.14, 0.41 |
| Weather:Sunny | 0.08 | 0.18 | 0.23 | 0.63 |  |  |  |

| Dewlap display |  |  |  |  |  |  |  |
| --- | --- | --- | --- | --- | --- | --- | --- |
| | Estimate | SE | $\chi^2$ | p | Type-S error | p from permutation test | Bootstrapped 95% CI |
| Intercept | -4.39 | 0.11 |  |  |  |  |  |
| Male conspecific | 0.11 | 0.13 | 0.70 | 0.40 | 0.02 | 0.30 | -0.08, 0.38 |
| **Female conspecific** | **0.53** | **0.13** | **15.22** | **<0.01\*** | **~0** | **0** | **0.30, 0.80** |
| Weather:Sunny | 0.04 | 0.09 | 0.19 | 0.66 |  |  |  |
| **Patagium extension** |  |  |  |  |  |  |  |
| | Estimate | SE | $\chi^2$ | p | Type-S error | p from permutation test | Bootstrapped 95% CI |
| Intercept | -3.72 | 0.20 |  |  |  |  |  |
| **Male conspecific** | **0.68** | **0.21** | **8.35** | **0.002\*** | **~0** | **0** | **0.31, 1.23** |
| Female conspecific | 0.16 | 0.25 | 0.40 | 0.51 | 0.05 | 0.31 | -0.11, 0.47 |
| Weather:Sunny | 0.22 | 0.17 | 1.49 | 0.20 |  |  |  |

**Figure 5:**
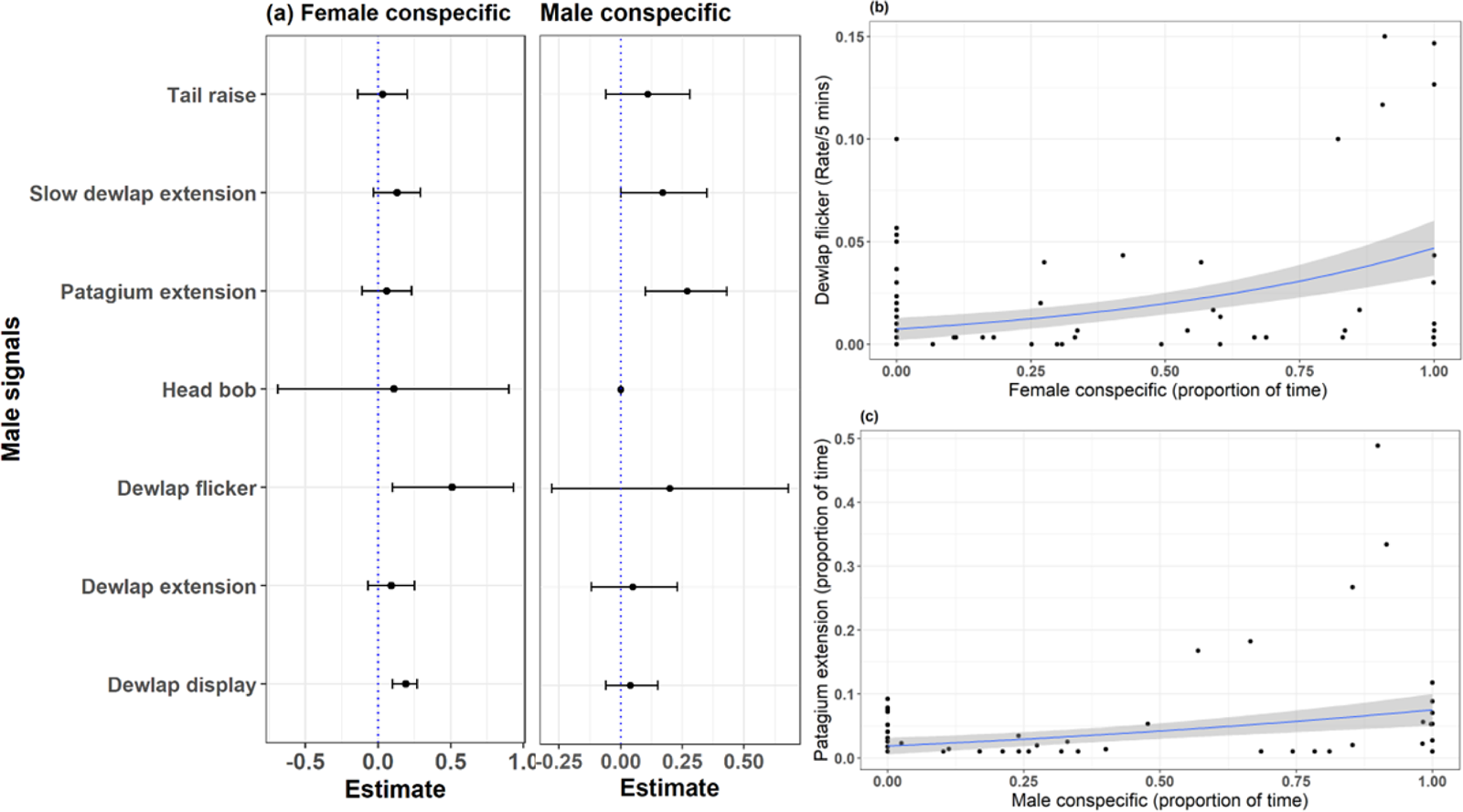
Association of male signals with the presence of male and female conspecifics. (a) Summary of standardised model predictions as represented by effect size and 95 percent confidence interval. (b) and (c) The model prediction represented by the regression line shows the relationship of male signals with female and male conspecifics. Dewlap flicker (b) was positively related to the presence of female conspecifics and patagium extension (c) was positively related to the presence of male conspecifics. Black dots represent actual data points.

## Discussion

We found that both female and male *Draco* lizards socially interact with conspecifics using a variety of signals. An interesting feature of the study is the diverse set of signals observed in females. Both males and females deployed similar visual signals with remarkable use of the dewlap. Examining the social context in which these signals were employed, both females and males appear to be using them to interact with conspecifics. Furthermore, these signals seem to be maintained by both the mechanisms of multiple receiver and backup signal hypothesis.

### Maintenance of multiple signals in the sexes

Both sexes in *D. dussumieri* employ a striking set of visual signals involving their dewlap, patagium and body postures to engage with both male and female conspecifics. In females, the relationship of signals with social context showed a clear support for the backup signal hypothesis and a relatively weak support for the multiple receiver hypothesis. A subset of the multiple signals (e.g., patagium extension and arched back) were correlated (Figure 2) indicating support for the backup signal hypothesis (Moller & Pomiankowski, 1993). These correlated signals may therefore be maintained to reinforce the information being conveyed through the signals (Candolin, 2003). Studies examining the maintenance of multiple signals in females are relatively few. Recent work on birds reports a positive correlation between acoustic and visual trait elaboration in females across multiple species, implying support for the back-up signal hypothesis (Webb et al., 2016; Jones et al., 2021). Our analysis also suggested an association between female signals like dewlap display and patagium extension with the presence of conspecific females and males respectively, suggesting support for the multiple receiver hypothesis (Johnstone, 1997b), but the effect size for some of the signals was variable and needs further investigation (Table 1). The association of signals in female *Draco* with the presence of other female conspecifics suggests the use of these signals in intra-sexual competition (Reedy et al., 2017; Ranade et al., 2022) and warrants further investigation.

The multiple signals in male *Draco* lizards were also utilised to interact with both sexes and seem to be maintained by both mechanisms (Table 2). Signals like dewlap flicker and dewlap display were primarily performed in the presence of conspecific females, while patagium extension was done mainly in the presence of conspecific males. Male fan-throated lizards, *Sarada superba*, have been reported to exhibit a comparable pattern where distinct signals elicit responses from different audiences (Zambre & Thaker, 2017). These multiple signals in male *Draco* may therefore be maintained as different receivers pay attention to different signals (Andersson et al., 2002; Candolin, 2003) or the different signals reflect different aspects of the individual’s quality (Martín & López, 2009). There was also some support for the backup signal hypothesis, as a few signals like dewlap extension and slow dewlap extension were correlated (Figure 3).

Moreover, several signals in both sexes were associated neither with female nor male conspecifics in the vicinity, suggesting that these signals might be utilised in some other social contexts and can be investigated in future studies. Overall, both the backup and multiple receivers mechanisms seem to be involved in maintaining multiple signals in both females and males. Other studies (e.g., Eland antelope *Tragelaphus oryx*, Bro-Jørgensen & Dabelsteen, 2008) similarly report that multiple mechanisms may be important in the maintenance of multiple signals within populations.

### Sex difference in signalling behaviour

Consistent with the accumulating evidence for the prevalence of elaborate traits and behaviours in females, the signalling repertoire of female *Draco* was as diverse as that of males. Males and females, however, differed in the relative use of those signals. Some signals (e.g., arched back, patagium extension) were used more commonly by females; whereas most dewlap-related signals were performed more by the males (Supplementary Table 2). Similar findings have been recorded in *Anolis* lizards, where the signalling repertoire was similar between the sexes; however, they differed in their use (Jenssen et al., 2000; Harrison & Poe, 2012; Driessens et al., 2014, 2015). This variation in the use of signals might indicate either a difference in the costs and benefits associated with signalling (T. Clutton-Brock, 2007; Cain & Ketterson, 2013) or the use of these signals in specific contexts (Peters et al., 2016; Jones et al., 2022).

To infer the potential function of signals, we investigated the association of signals with two social contexts, i.e., intra-sexual and inter-sexual interactions. Both sexes utilised signals to interact with conspecifics in both social contexts. However, some signals like dewlap display were used in different social contexts by each sex, indicating different functions of those signals in the sexes. Patagium extension was used by males only in same-sex interactions, but females seemed to be utilising it in both social contexts. These findings are similar to those observed in birds where plumage (Bennett et al., 1997; Swaddle & Witter, 1997), bill colouration (Johnson et al., 1993; Murphy et al., 2009) and songs (Mason et al., 2014; Webb et al., 2016) often serve different functions and are maintained by different mechanisms in the sexes. This divergence could be an outcome of the different selective pressures acting on the sexes as a result of their distinct life histories and reproductive roles (Tobias et al., 2012; Dale et al., 2015; Odom et al., 2016; Wilkins et al., 2020; Odom et al., 2021; Enbody et al., 2022). These studies also highlight that contrary to conventional understanding, sexual dimorphism is not solely an outcome of sexual selection in males. Rather, males and females are often under independent selection pressures, potentially resulting in signalling traits with distinct structure, function, and evolution (LeBas, 2006; Tobias et al., 2012; Price, 2015; Jordan Price, 2019; Enbody et al., 2022). Consequently, a better understanding of trait elaboration will be possible through a sex-inclusive research method that looks at elaborate characteristics and behaviours in both sexes (Riebel et al., 2019).

## Conclusion

Elaborate traits, ornaments, and display behaviour have traditionally been regarded as male-exclusive. Nonetheless, over the last decade, there has been increased documentation of similarly elaborate and competitive traits in females as well (T. Clutton-Brock, 2007, 2009; Rosvall, 2011; Stockley & Bro-Jørgensen, 2011; Tobias et al., 2012; T. H. Clutton-Brock & Huchard, 2013; Stockley & Campbell, 2013). In contrast to previous studies on the social behaviour of *D. dusumieri*, which described females as passive (John, 1962, 1966), the findings of this study document the use of multiple signals by females in different social contexts. The multiple signals in females were as diverse as those in males and could potentially serve a role in communication. While additional experimental investigation is necessary to validate the results, our findings highlight diverse signals in females, which may have different functions from those of males. Our study, hence, supports the need for sex-inclusive research for a better understanding of complex traits and behaviours (Riebel et al., 2019; Wilkins et al., 2020).

## Acknowledgements

We thank the National Centre for Biological Sciences for institutional and financial support. Nature Conservation Foundation and Society for Study of Amphibians and Reptiles (SSAR) for funding support. Kalinga Centre for Rainforest Ecology provided logistical support in the field. We thank the Wildlife Office, Dr Jayashree Ratnam, Dr Vivek Ramachandran and Alissa Barnes for administrative support. We are grateful to Daimler Pereira, Chinmayee and Nithin Veeresh for the immense help with the fieldwork.

## Supplementary

**Table 1:** Description of behaviours shown by both sexes. Signals are represented in bold.

| Behaviour | Description |
| --- | --- |
| Flat body | Body completely flat on the tree |
| Only head raised | Head raised, forelimb slightly raised and hind limbs flat with ventral part of the body in contact with tree |
| Forelimbs raised | Head raised, forelimb straightened and hind limbs flat with only hind part of the ventral body in contact with tree |
| All limbs raised | All four limbs raised and ventral part raised in air |
| **Tail raise** | The tail is raised from the tree surface |
| **Tail flick** | Side-to-side movement of the tail |
| Head movement | Rotation of head only- up/down/left/right |
| **Head bob** | Up-down movement of the head only |
| Shift | Shifting the body either left or right while being at same height |
| Move | Change in the vertical position on the tree (move up or down) |
| Forage | Feed on an ant/ insect |
| **Dewlap extension** | Complete extension and retraction of dewlap |
| **Slow dewlap extension** | The dewlap is extended and retracted at a slower speed |
| **Dewlap display** | Dewlap kept in an extended position for some time |
| **Dewlap flicker** | Flickering of dewlap in complete or semi-stretched extension |
| **Arched back** | The back is raised and bent upwards like an arch |
| **Patagium extension** | Complete extension of patagium |
| Glide | Glide from one tree to another |
| **Turn and circle** | Turn and then circle around another individual with complete tail raised and dewlap being displayed |
| Approach a male | Move towards a male |
| Approach a female | Move towards a female |
| Approached by male | Another male moves towards the focal individual |
| Approached by female | Another female moves towards the focal individual |
| Chase | Move towards another individual and the other individual runs away |
| Chased by | Run away when another individual approaches |
| Scratch | Scratch the body using its limb |

**Table S2:** Variation in the use of signals by sex. * represents events (rate/5 minutes) and # represents states (proportion of time spent). Number of focal observations: Males= 114, Females= 55

| Behaviour | Female |  |  | Male |  |  |
| --- | --- | --- | --- | --- | --- | --- |
|  | Mean<br>(Rate per 5<br>minutes or<br>proportion<br>of time<br>spent) | 95% CI<br>(Rate per 5<br>minutes or<br>proportion of<br>time spent) | %<br>Observation<br>signal seen | Mean<br>(Rate per 5<br>minutes or<br>proportion of<br>time spent) | 95% CI<br>(Rate per 5<br>minutes or<br>proportion of<br>time spent) | %<br>Observation<br>signal seen |
| Dewlap extension* | 12.6 | 8.5-17.5 | 87.27 | 16.1 | 13.9-18.4 | 94.73 |
| Slow dewlap extension | 0.3 | 0.007-0.7 | 14.54 | 5.3 | 4.5-6.1 | 87.71 |
| Dewlap flicker* | 0.6 | 0.2-3.2 | 5.45 | 1.2 | 0.7-1.8 | 56.14 |
| Dewlap display<sup>#</sup> | 0.002 | 0.0002-0.006 | 12.72 | 0.005 | 0.003-0.007 | 29.81 |
| Head bob* | 0 | 0 | 0 | 0.2 | 0.1-0.3 | 17.54 |
| Tail raise<sup>#</sup> | 0.06 | 0.03-0.1 | 41.82 | 0.08 | 0.05-0.1 | 46.49 |
| Tail flick<sup>#</sup> | 0.002 | 0.00001-0.005 | 14.54 | 0 | 0 | 0 |
| Patagium extension<sup>#</sup> | 0.03 | 0.02-0.06 | 36.36 | 0.02 | 0.01-0.04 | 30.70 |
| Arched back<sup>#</sup> | 0.03 | 0.009-0.05 | 29.09 | 0.005 | 0.0004-0.01 | 7.01 |
| Turn and circle<sup>#</sup> | 0 | 0 | 0 | 0.001 | 0.0006-0.002 | 12.28 |

